# Heart spheroids from primary cells for studying cardiac development

**DOI:** 10.1101/2025.05.05.652113

**Authors:** Cindy Chang, Juan Xu, Shirley Chang, Guang Li

## Abstract

Organoids can be broadly classified based on their source: either from primary cells or from *in vitro* stem cell differentiation. Organoids derived from primary cells have been developed for various organs such as the liver, intestine, and colon, demonstrating significant value in disease modeling and drug screening. However, primary cell-derived heart organoids have not yet been reported. In this study, we developed heart organoid-like spheroids using murine atrial and ventricular cells. We demonstrated that these spheroids largely preserved the cell lineages present *in vivo*. We further evaluated their responses to treatment with growth factors. Subsequently, we fused atrial and ventricular spheroids to observe the migration patterns of different cell types. Then, we generated mixed spheroids composed of cells from different heart regions or organs and found that region- or organ-specific cells tended to cluster together. Notably, liver- and intestine-derived cells promoted cardiomyocyte maturation within these mixed organoids. Finally, we performed a fibroblast ablation assay in heart spheroids derived from transgenic mice and observed the effects on other cell lineages. Overall, we successfully generated heart spheroids from murine primary cells and demonstrated their *in vivo*-like characteristics. Compared to stem cell-derived organoid systems, this primary cell-derived approach holds great promise for translational research due to its potential to better preserve native cellular features.

## Introduction

Three-dimensional (3D) cellular aggregates, such as organoids and spheroids^1^, have become widely used due to their ability to model *in vivo*-like environments and cell interactions. These aggregate structures can develop from either pluripotent stem cells or tissue-derived primary cells^2^. Stem cell-derived organoids offer greater lineage plasticity^3^, while primary cell-derived organoids better preserve the genetic and cellular identities of the original tissue, making them complementary for many applications. Primary cell-derived organoids have been successfully developed for multiple tissues, including the liver, colon, intestine, breast, and various types of cancers^4,5^. These organoids have been extensively used to model normal physiological processes, disease progression, and for drug screening. However, primary tissue-derived heart organoids have not yet been developed.

The heart developmental processes in mice and humans are largely consistent^6^. At early stages, such as embryonic day 10.5 (E10.5) in mice, the heart consists of four chambers: the left ventricle (LV), right ventricle (RV), left atrium (LA), and right atrium (RA). In addition to these chambers, the heart contains two non-chamber structures: the atrioventricular canal (AVC), which is located between the atria and LV and later develops into the atrioventricular valves and septum; and the outflow tract (OFT), which connects to the RV and later forms the semilunar valves and major cardiac vessels such as the aorta and pulmonary artery^7,8^. These distinct heart regions originate from two progenitor sources known as the first heart field (FHF) and the second heart field (SHF). The FHF primarily contributes to the development of the LV, AVC, and portions of the atria, while the SHF gives rise to the RV, OFT, and parts of the atria^9,10^.

Heart development is a complex process regulated by numerous signaling pathways^11,12^. WNT signaling plays distinct roles at different stages of heart development: it is activated to specify mesodermal fate and subsequently repressed to allow cardiac mesoderm differentiation. Later, reactivation of WNT signaling in cardiomyocytes (CMs) promotes their proliferation^13,14^. VEGF signaling is another critical pathway that primarily promotes the development of endothelial cells (ECs). The heart contains two main types of endothelial cells that both require VEGF for proper development^15,16^: endocardial endothelial cells (EndoECs) and vascular endothelial cells (Vas_ECs). Retinoic acid (RA) is a multifunctional molecule; its signaling pathway is essential for atrial CM lineage specification during early development^17,18^. Additionally, RA has been shown to promote ventricular CM maturation^19^ and regulate the topological organization of brain organoids^20^.

Heart development and function is also regulated by interactions with other organs. At early embryonic stages, such as E14.5, major visible organs include the heart, lung, liver, kidney, stomach, intestine, and brain^21^. The lung and liver, located adjacent to the heart, contribute cell lineages and growth factors to each other, influencing their mutual development. The kidney and heart, both derived from the mesoderm layer, function closely together within the circulatory system. Foregut tissues, including the liver, stomach, and intestine, also co-develop with the heart and engage in active signaling crosstalk^22^. The brain connects to the heart through the nervous system, regulating not only heart contractions but also heart growth via signaling molecules. Since stem cell-derived heart organoids often contain non-cardiac lineages and multi-lineage organoids have previously been shown to promote cardiac lineage development^23,24^, it is important to directly assess the interactions between cardiac cells and cells from these associated organs.

In addition to cardiomyocytes (CMs), the heart contains many other cell types, such as fibroblasts. To study their role in heart development, we employed a conditional ablation system based on Diphtheria Toxin fragment A (DTA)^25^. DTA, a gene-encoded cytotoxin, has been widely used to selectively eliminate target cells in various tissues. In Rosa26-DTA mice, DTA expression is controlled by Cre recombinase-mediated gene recombination. To specifically ablate fibroblasts, we crossed Rosa26-DTA mice^26^ with fibroblast-specific Pdgfra-CreER mice^27^ and administered tamoxifen to pregnant females to induce cell ablation at desired embryonic stages. However, since *in vivo* ablation affected fibroblast populations and organ development across multiple tissues, the heart defects observed could have been influenced by defects in other organs. Therefore, it would be advantageous to assess the role of fibroblasts in cardiac development using a heart-specific *in vitro* or *ex vivo* system.

In this study, we developed a heart spheroid system using primary mouse heart cells. We demonstrated that these spheroids preserved key cell lineages and identities. Additionally, we investigated their responses to growth factors and their interactions with cells from other organs. Finally, we demonstrated the system’s feasibility in modeling deficiencies caused by the perturbation of specific cell types. This heart spheroid platform holds significant potential for basic research in heart development, disease modeling, and drug screening in both mouse and human systems.

## Results

### Development of heart spheroids from murine primary cells

To generate heart spheroids, we isolated mouse primary heart cells from early embryonic stages and aggregated them in ultra-low attached 96-well plates. We seeded a range of cell densities per well ranging from 2.5k to 100k and tested two culture mediums, including a DMEM based and an IMDM-based medium (Fig S1A, details in the experimental methods section). After analyzing the spheroids at day 4, 7, and 12, we observed that spheroids generated from 10k cells cultured in IMDM-based medium have round morphologies and lack dark areas in the middle, an indicator of extensive cell death, confirming their suitability for subsequent experiments. Using this condition, we generated spheroids from atrial and ventricular cardiac cells at embryonic day 10.5 (E10.5) and E14.5. We observed that at both stages, atrial spheroids have around 140 beats per minute, while ventricular spheroids have less than 50 beats per minute (Fig 1A, B). Next, we performed immunofluorescence (IF) analysis of cardiac lineage gene markers in the spheroids. First, we analyzed the expression of atrial and ventricular CM lineage genes. In atrial spheroids, we observed the expression of the general CM marker MF20 and atrial-specific CM lineage marker Myl7, while in ventricular spheroids, we observed the expression of general CM marker Troponin and ventricular CM lineage-specific marker Myl2 (Fig 1C). We also confirmed the identity of atrial CMs in atrial spheroids using another atrial marker gene Nr2f2 and observed similar results (Fig S2A). Moreover, we analyzed the expression of genes for different cell lineages. We found that the MF20+ CM were largely evenly distributed in the spheroids, while the Vim+ fibroblasts were mainly located on the edges. We also analyzed the endothelial cell marker Cd31 and EndoEC marker Nfatc1. We found that most Cd31+ cells were EndoEC and mainly located in the middle of the spheroids (Fig 1D).

**Figure 1.**
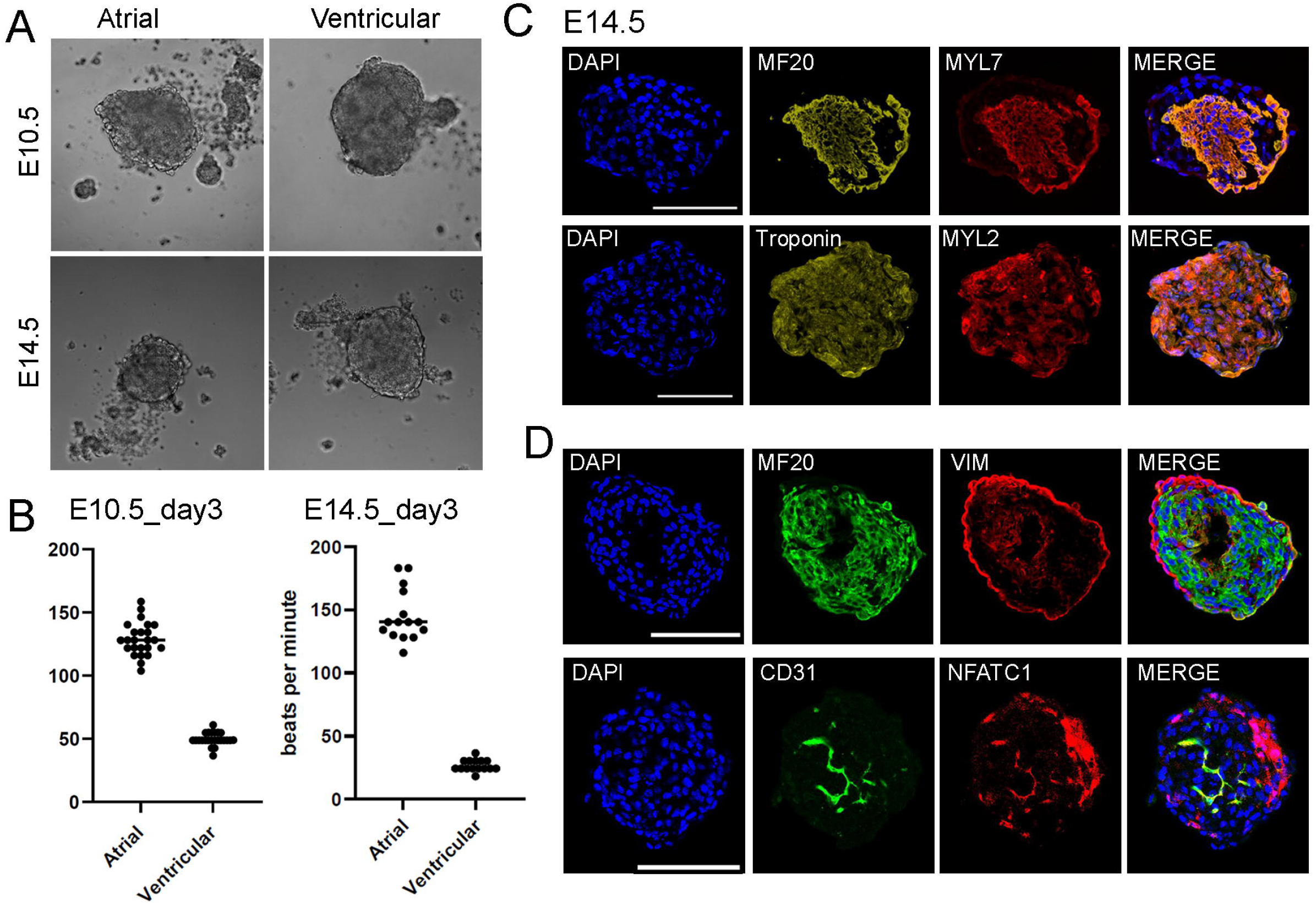
Primary mouse cell-derived cardiac spheroids recapitulate *in vivo* like embryonic heart features. (A) Representative brightfield images of atrial and ventricular spheroids generated with e10.5 and e14.5 mouse heart cells. Scale bar=100 μm. (B) Beating rates of atrial and ventricular spheroids on day 3. (C, D) Immunofluorescence images of e14.5 atrial and ventricular spheroids staining for cardiac cell lineage genes. Scale bar=100 μm.

### Growth factor treatment of the heart spheroids

Next, we tested the response of the spheroids to three growth factors that are important for cardiac development. We respectively treated E10.5 atrial and ventricular spheroids with WNT agonist CHIR, VEGF, and retinoic acid (RA) (Fig 2A). We observed that CHIR treatment led to reduced beating rates in both atrial and ventricular spheroids, while VEGF treatment increased the beating rate in atrial spheroids but not in ventricular spheroid, and RA treatment did not result in significant changes in both types of spheroids (Fig 2B). Quantification of the spheroid size showed that CHIR treatment led to larger spheroids in both atrial and ventricular spheroids, while VEGF and RA treatments did not lead to significant size changes in comparison to the control (Fig 2D, E). Subsequently, we conducted IF staining for MF20 and Cd31 (Fig 2C). We observed that Cd31 signal was reduced after CHIR treatment and increased after VEGF treatment in both atrial and ventricular spheroids. Interestingly, RA treatment also led to a decrease of Cd31 positive area in ventricular spheroids but not in atrial spheroids. For MF20 staining, both CHIR and VEGF treatments led to a decrease of MF20+ CM areas in both atrial and ventricular spheroids. Lastly, we analyzed cell proliferation in the spheroids by staining for pHH3. We observed that only CHIR treatment significantly increased cell proliferation (Fig 2D, E). These results suggest that the spheroids respond differently to the growth factor treatments.

**Figure 2.**
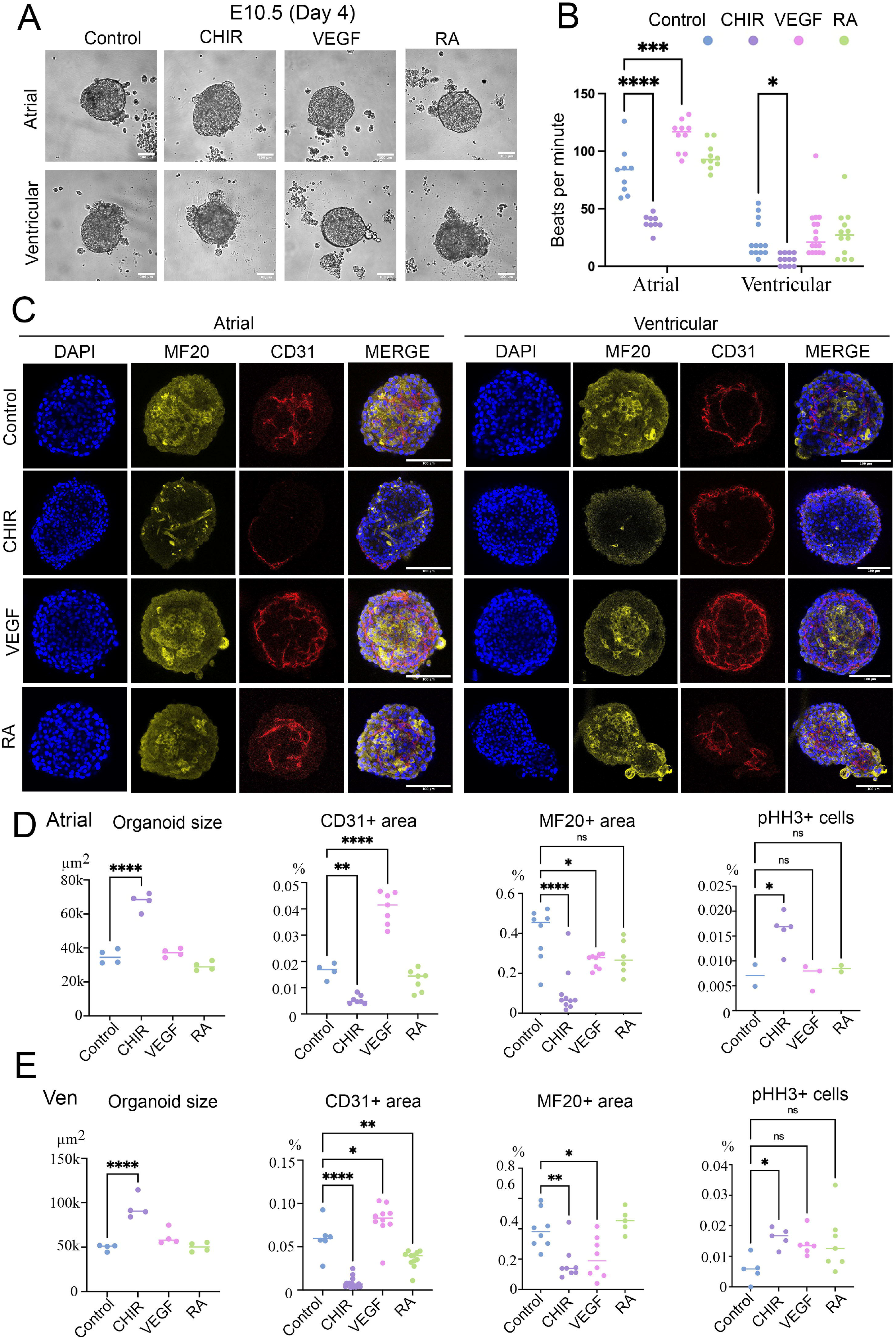
Modulation of spheroid development with growth factors treatment. (A) Representative brightfield images of atrial and ventricular spheroids treated with different growth factors on day 4. Scale bar=100 μm. (B) Beating rates of the spheroids treated with different factors on day 4. (C) Immunofluorescence images of atrial and ventricular spheroids treated with different growth factors staining for CM and endothelial cell lineage genes. Scale bar=100 μm. (D, E) Quantification of spheroid size, EC area, CM area, and pHH3+ cells in atrial and ventricular spheroids with the treatment of different growth factors. * represents p < 0.05, ** represents p < 0.01, **** represents p < 0.001.

### Fusion of spheroids from different heart regions

Afterward, to assess if additional heart structures such as the atrioventricular canal (AVC) can be induced in our spheroid systems, we fused one atrial spheroid and one ventricular spheroid. To track the movement of cells from individual spheroids, we fused one GFP+ spheroid which has atrial or ventricular identity with one GFP-spheroid which has ventricular or atrial identity (Fig 3A). In 7 days, however, we failed to observe the induction of extra structures at the fusion site. In contrast, we observed that GFP+ and GFP-cells were still largely spatially distinct. We further conducted IF staining to analyze the identity of the GFP+ cells (Fig 3B). Interestingly, we found that CMs remained in their original spheroids with minimal migration, fibroblasts (using marker Col1a1) displayed subtle migration along the organoid surface, and EC was the major cell type that migrated across the fusion boundary (Fig 3C). We further quantified their beating rates and observed that fused spheroids had similar rates to atrial spheroids (Fig 3D). Lastly, we generated spheroids by mixing GFP+ atrial cells with RFP+ ventricular cells in a 1:1 ratio. Interestingly, we observed that the GFP+ and RFP+ cell populations were largely distinct within the mixed spheroids (Fig 3E). This result suggested that the atrial and ventricular cells preferably aggregate together with cells of the same lineages. Additionally, we generated spheroids for the six cardiac sub regions in early embryonic stages (E10.5) including LA, RA, AVC, LV, RV, and OFT (Fig S3A, B). We measured their beating rates at 4 days post aggregation and observed relatively high beating rates in the LA, RA, and LV-derived spheroids, and lower beating rates in OFT, RV, and AVC-derived spheroids (Fig S3C). Furthermore, we fused spheroids derived from adjacent cardiac regions, pairing one spheroid labeled RFP+ with another labeled RFP-.Consistent with atrial and ventricular fused spheroids, we did not observe the induction of extra heart structures and the cell populations of each cardiac region remained largely separated (Fig S3D).

**Figure 3.**
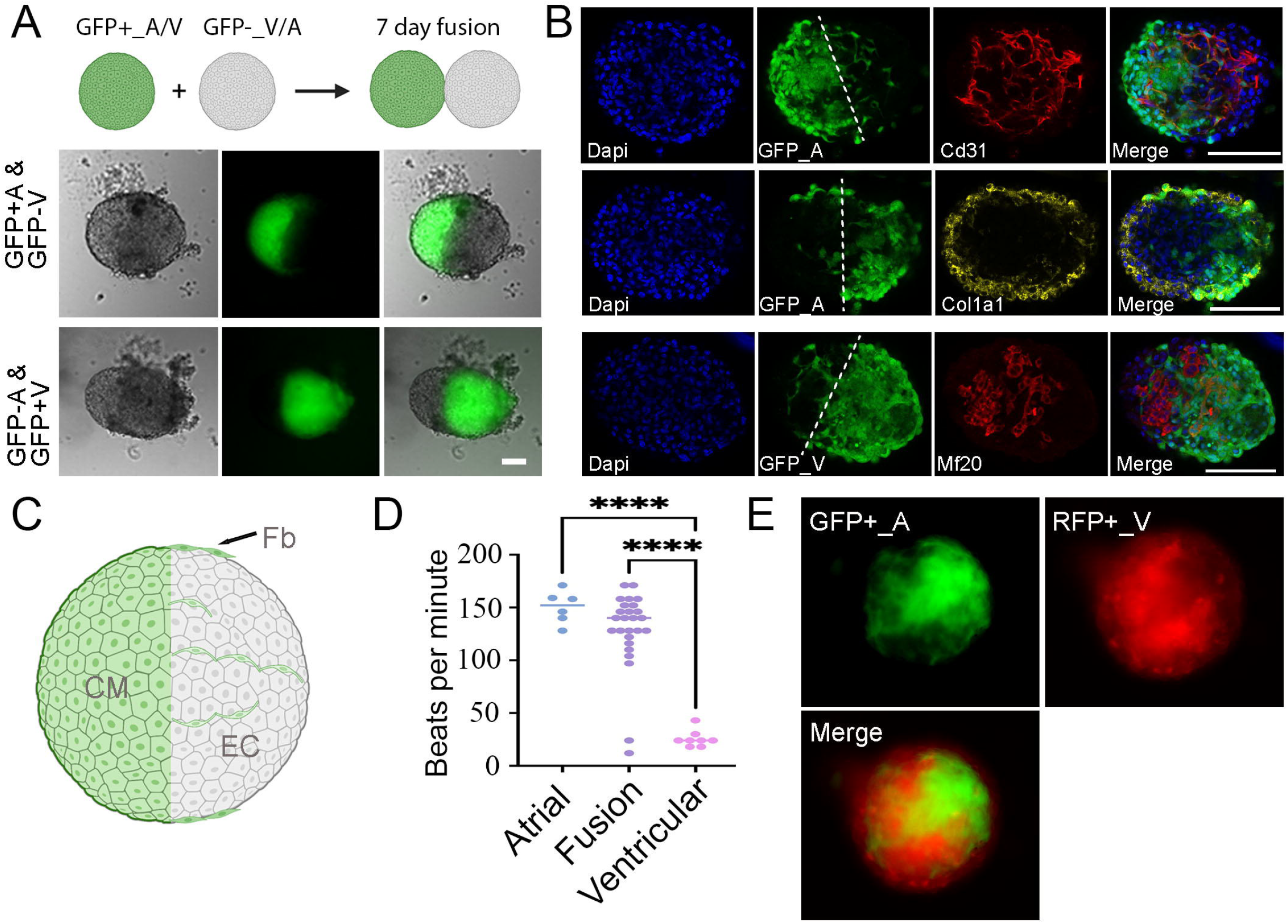
Cell migration in atrial and ventricular fused and mixed spheroids. (A) Representative brightfield and fluorescent images of e10.5 atrial-ventricular fused spheroids after 7 days of fusion. Scale bar=100 μm. (B) IF images of fused spheroids staining for different lineage genes. Scale bar=100 μm. (C) Schematic of cell migrations in the fused spheroid. (D) Beating rates of e10.5 fused spheroids on day 2. (E) Fluorescent images of GFP+ atrial cells and RFP+ ventricular cells in mixed spheroids.

### Interaction analysis of cells from different tissues in mixed spheroids

We then analyzed the impact of other organ cells on cardiac cell development by generating mixed spheroids. We generated the mixed spheroids with a 3:1 ratio of heart cells (atrial or ventricular) and cells from other organs including brain, lung, kidney, liver, stomach, and intestine. We selected these organs because they are physically adjacent to the heart *in vivo* or frequently co-developed with cardiac lineages in stem cell-derived organoids. We labeled cardiac cells with RFP and other tissue cells with GFP. Interestingly, we found that non-cardiac tissue cells were spatially distinct from the cardiac cells, with brain and kidney-mixed spheroids having the most obvious separation (Fig S4). Next, we quantified the beating rates of the spheroids. We observed that atrial spheroids mixed with brain, stomach, or intestine cells have significantly lower beating rates than the atrial spheroid controls, while ventricular spheroids mixed with brain, kidney, or intestine cells have lower beating rates than their control counterpart (Fig 4B). Moreover, we analyzed CM maturation by quantifying the ratio of *Myh6* to *Myh7* expression with qPCR. We observed that atrial spheroids mixed with liver or intestine cells had a higher *Myh6/Myh7* ratio than the atrial control, indicating an improved maturation in these mixing spheroids. In contrast, we observed that ventricular spheroids mixed with brain, lung, liver, or intestine cells were more mature than ventricular spheroid controls (Fig 4C). These results indicate that co-culture with cells from certain but not all organs can improve CM maturation.

**Figure 4.**
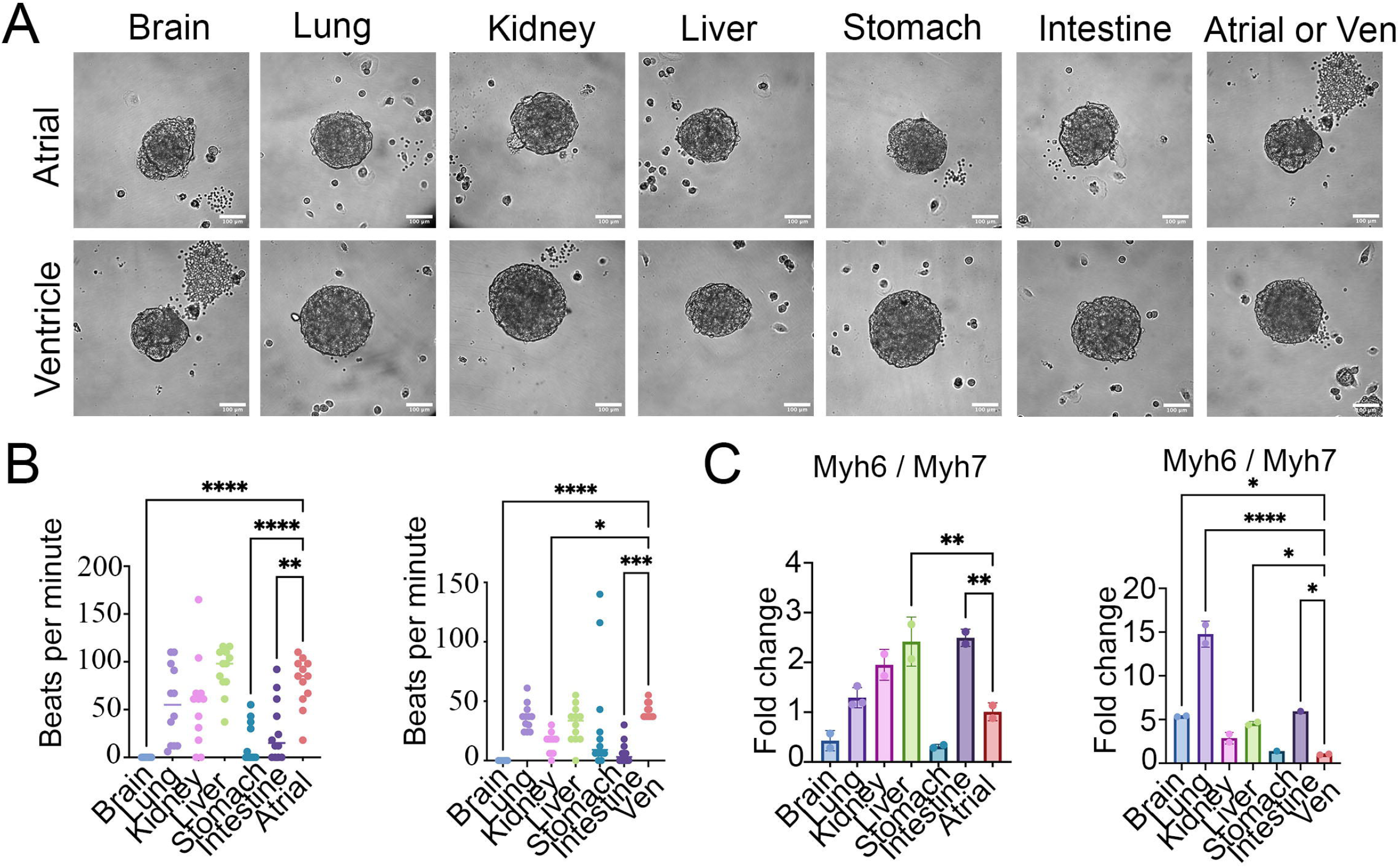
Functional analysis of other organ-derived cells in cardiomyocyte development. (A) Representative brightfield images of spheroids composed of e14.5 atrial or ventricular cells and other organ-derived cells. Scale bar=100 μm. (B) Beating rates of the mixed spheroids on day 6. (C) *Myh6* to *Myh7* gene expression ratio of spheroids mixed with different organ cells. * represents p < 0.05, ** represents p < 0.01, **** represents p < 0.001.

### Modeling of cell ablation assay in heart spheroids

Subsequently, we generated ventricular spheroids from Pdgfra-CreER+/-; Rosa-DTA+/-mice and Rosa-DTA+/-mice hearts. In Pdgfra-CreER+/-; Rosa-DTA+/-spheroids, the fibroblast were ablated after 4-HT treatment and used as an experimental group. In contrast, because Rosa-DTA+/-spheroids will not respond to 4-HT treatment, they served as controls (Fig 5A). We treated both control and experimental spheroids with three different concentrations of 4-HT including 0 μM, 0.5 μM, and 2 μM. We found that spheroid sizes in the experimental groups treated with 0.5 μM and 2 μM 4-HT were significantly reduced compared to the control groups treated with the same concentrations. A similar reduction was observed when compared to the experimental group treated with 0 μM 4-HT (Fig 5A). Next, we conducted IF staining for Caspase 3 to analyze cell death in the spheroids. However, we did not observe a significant increase in Caspase 3 signal (Fig. 5C), likely because most of the ablated cells had already died at this stage. We further quantified the EC-positive area in the ablated spheroids (Fig. 5B). We observed that the 4-HT treated spheroids had a reduced percentage of EC+ area in the control group, indicating potential side effects from 4-HT treatment. In contrast, we observed that in the experimental group, 4-HT treatment increased the percentage of EC+ area. We also analyzed cell proliferation in the spheroids by staining for EdU and observed a similar trend to the Cd31 staining results (Fig 5C). These results indicate that fibroblast ablation could have a positive impact on cell proliferation and EC growth in the spheroids.

**Figure 5.**
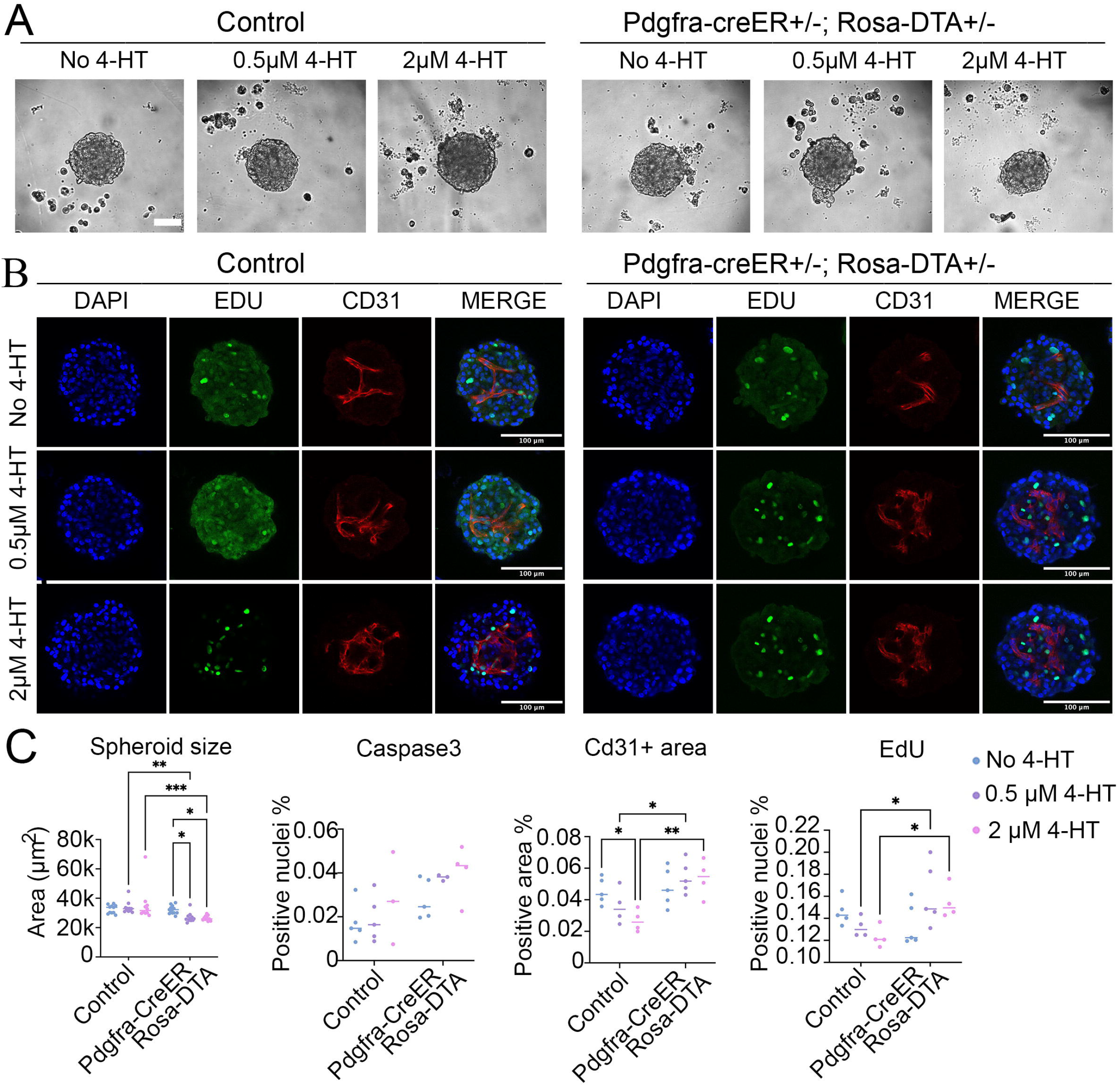
Modeling heart developmental deficiency in Pdgfra-CreER+/-; Rosa-DTA+/-mouse heart-derived spheroids. (A) Representative brightfield images of control and experimental Pdgfra-CreER+/-; Rosa-DTA +/-ventricular spheroids with different 4-HT concentrations. (B) IF images of control and ablated spheroids for EC lineage gene Cd31 and cell proliferation marker EdU. Scale bar=100 μm. (C) Quantification of spheroid size, Caspase 3 signal, Cd31+ cell area, and Edu signal in control and ablated spheroids with different 4-HT concentrations.

## Discussion

In this study, we generated spheroids from different regions of the heart and examined their responses to three growth factors. We also fused spheroids from distinct regions and observed cell type-specific migration patterns. Additionally, we created mixed spheroids containing cardiac cells and cells derived from other organs, finding that some non-cardiac organ-derived cells promoted cardiomyocyte maturation. Finally, we performed fibroblast ablation experiments using the spheroid system to demonstrate its potential for dissecting the roles of specific cell types in heart development. This system also shows promise for modeling heart diseases using human patient-derived cardiac cells.

Unlike stem cell-derived organoids, organoids derived from primary cells can preserve key characteristics of their *in vivo* counterparts, providing a more accurate representation of cardiac lineages, their relative proportions, and the maturation status of each cell type. Additionally, for certain rare human patient samples where stem cell-based models are not yet available, generating spheroids from primary cells offer a valuable approach to preserving these samples and studying their pathology. Moreover, while stem cell-derived organoid systems have not been developed for most mouse organs, and numerous transgenic mouse lines exist that carry specific mutations, fluorescent reporters, and lineage-tracing tools, the primary cell-derived spheroid system can leverage these resources to investigate fundamental processes in organ development and physiology. These analyses are essential, as they can complement similar studies in human organoids derived from primary tissues.

In this study, we treated cardiac spheroids with three growth factors that are essential for the proper development of the heart. Activation of the Wnt pathway promoted spheroid growth but compromised cellular identity, leading to increased proliferation at the expense of chamber-specific marker expression. VEGF, a key angiogenic factor, enhanced vascularization but caused a slight reduction in the cardiomyocyte region. In contrast, RA treatment had minimal effects on overall spheroid growth or beating rates, but specifically reduced the endothelial cell region in ventricular spheroids, suggesting a potential role in modulating region-specific angiogenesis.

These findings highlight how different growth factors influence organoid development and maturation, offering insights into the roles of signaling pathways in cardiac development. Additionally, this knowledge can guide improvements to the spheroid system, for instance, using Wnt and VEGF to respectively enhance growth and vascularization. Moving forward, the incorporation of growth factors from additional signaling pathways may facilitate passaging and long-term maintenance of cardiac spheroids, further enhancing the utility of this system.

We fused spheroids derived from different heart regions but did not observe the formation of any new major structures. Given that the cardiac cells used were from the early embryonic stage (E10.5) and should retain certain cellular plasticity, the absence of new structures such as the AVC following fusion of atrial and ventricular spheroids may be due to disrupted signaling gradients or the absence of key extracellular cues. These results suggest that the inclusion of the correct cell types may not be sufficient to form the expected structures, raising caution when testing similar hypotheses with spheroids generated using other technologies, such as bioprinting.

Additionally, when atrial and ventricular cells were mixed within the same spheroid, we observed that cells from the same chamber tended to cluster together, suggesting that lineage specific cells possess mechanisms to recognize and aggregate with one another. Identifying the molecular pathways underlying this behavior will be an important direction for future research. Interestingly, similar aggregation behavior have been observed in stem cell-derived heart organoids, where atrial and ventricular cardiomyocytes also preferentially cluster together^28^, indicating this may be a conserved property shared between mouse primary cells and human iPSC-derived cells.

Furthermore, when cardiac cells were mixed with cells from other organs, we observed a similar tendency for cells to self-segregate by tissue type. Notably, each combination of mixed tissues displayed unique spatial patterns, raising intriguing questions about the mechanisms driving these arrangements and their implications for interorgan interactions.

## Methods

### Experimental Methods

#### Mouse strains

All animal procedures were conducted in accordance with protocols approved by the Institutional Animal Care and Use Committee (IACUC) at the University of Pittsburgh. CD1 male and female mice (Charles River Laboratories) were housed and bred in-house to obtain embryos at defined developmental stages for spheroid generation. Transgenic mouse lines used in this study included Rosa26-mTmG (Strain #:007676)^29^, Pdgfra-CreERT2 (Strain No. 032770)^27^, and ROSA26-eGFP-DTA (Strain No. 032087) ^26^, all of which were obtained from The Jackson Laboratory.

#### Primary tissue dissection and spheroid culture

Unless stated otherwise, CD1 mouse strain was utilized to culture the cardiac spheroids. The mouse dissection and heart cell preparation were conducted following a previously published procedure^25^. Specifically, after dissection, all tissue samples were suspended in 1x PBS over ice in microcentrifuge tubes. Surgical scissors were used to manually cut the tissues into smaller pieces and then centrifuged at 300g for 3 minutes before resuspension in 300ul 0.25% Trypsin/EDTA (Gibco, 25200056). The tubes were incubated in a 37°C water bath for 15 minutes with incremented mixing with a P1000 pipette to keep the samples suspended. 300ul 20 mg/mL collagenase A and B (Sigma, 10103578001, 11088807001) was then added to the tubes and incubated for another 15 minutes in a 37°C water bath. After the incubation period, 600ul mouse differentiation media was added to the tubes. For the initial testing experiments, DMEM based medium (DMEM, 10% FBS, 2% Pen/Strep) and IMDM based medium^30^ (82% (v/v) IMDM, 2 mM L-glutamin, 15% (v/v) serum, 0.01‰ (v/v) monothioglycerol and 50 µg/ml ascorbic acid) were used, and the following experiments only used the IMDM-based medium. The cells were then filtered through Corning 100 Micrometer Cell Strainers (Corning, 431752) into new microcentrifuge tubes and then centrifuged for 5 minutes at 1000rpm at 4°C before resuspension in 1mL mouse differentiation media.

The primary cells were then counted on Nexcelom Bioscience Cellometer Auto 2000 and seeded at 10,000 cells/well in round bottom ultra-low attachment 96-well plates (Fisher, 07201680) at a volume of 200ul per well. The plate was centrifuged at 1000rpm for 10 minutes and then placed into an incubator. Half of the media was changed every other day until the spheroids were ready for analysis.

For experiments that were co-cultured with non-cardiac tissues, E14.5 mice were used and a 3:1 cardiac-other tissues ratio was utilized to plate 10,000 cells/well. For the spheroid fusion and mixing experiments, the RFP spheroids were generated using cardiac cells from the Rosa26-mTmG mice, while the eGFP spheroids were created using cells from the ROSA26-eGFP-DTA mice, which do not express DTA when the mice were not bred with Cre mice.

#### Growth factor and tamoxifen (4HT) treatment

E10.5 CD1 cardiac mice tissue were utilized in growth factor treated experiments. After cell differentiation and cell counting, atrial and ventricular samples were thoroughly mixed and then separated into four groups of equal volume: a negative control group, CHIR99021 (Selleck Chemical LLC, S292425MG), VEGF (GeminiBio, 300-827P), and RA (Sigma, R2625-50MG) treated groups. Treatment samples were centrifuged for 5 minutes at 1000rpm and 4°C and the supernatant was removed. New media was prepared with the different growth factors. Mouse differentiation media with a final concentration of 2 μM CHIR99021 was prepared and used to resuspend the cells in the CHIR99021 treated group. Similarly, VEGF and RA were added to their respective groups with final concentrations of 100ng/ml and 1 μM, respectively. The resuspended cells were plated 10,000 cells/well in round bottom ultra-low attachment 96-well plates and treated with appropriate concentrations of CHIR99021, VEGF, and RA. Fresh media was prepared and was changed every other day until 10 days of culture.

E17.5 Pdgfra-CreER +/-;Rosa26-DTA-eGFP+/-mice ventricular tissue were utilized for experiments treated with 4HT (Sigma, H7904-5MG). Cells were dissociated as described in the previous section and the ventricles of each embryo of a single litter was plated individually in mouse differentiation media. On day 3 of culture, half of the media was removed and the spheroids were divided into three groups per embryo, including a negative control group. On the initial day of treatment, one group was treated with a 4HT concentration of 1 μM and another was treated with 4 μM of 4HT to create final concentrations of 0.5 μM and 2 μM 4HT, respectively. New 4HT was prepared and the media was changed everyday for 5 days of treatment.

#### Live imaging and heartbeat tracking

Bright-field live imaging of the spheroids was captured using a Leica microscope. To calculate the average heart beating rate of the spheroids, videos of each spheroid were recorded and analyzed afterward.

#### Whole-mount organoid immunofluorescence staining and imaging

Spheroids were transferred to microcentrifuge tubes using a 1000ul pipette tip and fixed in 4% paraformaldehyde solution overnight at 4°C. The following day, the spheroids were washed 3 times in 1xPBS (5 minutes each) and 3 times (10 minutes each) in blocking buffer (PBS, 10% FBS, 0.2% Triton X-100) in room temperature. They were then blocked for 1 hour at 4°C in blocking buffer before being incubated overnight with primary antibodies in blocking buffer overnight at 4°C while shaking. The next day, the samples were washed 4 times (20 minutes each) with blocking buffer at 4°C while shaking before being incubated overnight with secondary antibodies and DAPI (1mg ml-1, Sigma-Aldrich) in blocking buffer overnight at 4°C while shaking. On the final day, the samples were washed for 1 hour with blocking buffer at 4°C while shaking and then rinsed in rinse buffer (PBS, 0.2% FBS, 0.2% Triton X-100).

For imaging, the samples were mounted on slides and imaged using Leica confocal laser scanning microscopy. Images were analyzed using Fiji (https://imagej.net/Fiji). DAPI positive cells were counted and used for normalization against the target cell marker of interest for cell quantification in the spheroids.

#### RNA extraction and qRT-PCR

Spheroids were collected in TRIzol (Invitrogen, 15596026) and RNA was extracted using the RNeasy Micro kit (Qiagen, 74004) according to manufacturer’s instructions. RNA was reverse-transcribed with the iScript cDNA Supermix kit (Biorad, 1708891). 3ul of cDNA per reaction was used for a total volume of 20ul. qPCR was run with a BioRad CFX Connect Real-Time PCR Detection System, using Power SYBR Green PCR Master Mix (Applied Biosystems, A46109), with an annealing temperature of 60°C. Gene expression was normalized to *Myh7* expression and relative fold expression was calculated with the 2-∆∆CT method.

#### EdU Cell Proliferation Assay

EdU Cell Proliferation Assay (Thermo Fisher, C10337) was performed following the manufacturer’s protocol. Briefly, a 2X working solution of EdU was prepared in mouse differentiation media from the 10mM stock solution (10 μM final concentration) and added in equal volume to the plates when changing media (half of old media removed at a time). The plates were incubated overnight in a 37°C incubator and then collected in labelled microcentrifuge tubes using a 1000ul pipette tip. The samples were then fixed in 4% PFA in PBS and incubated for 15 minutes at room temperature before being washed twice with 1mL of 3% BSA in PBS. After washing, 1mL of 0.5% Triton X-100 in PBS was added to each microcentrifuge tube and incubated at room temperature for 20 minutes. The permeabilization buffer was removed and the samples were washed twice with 1mL of 3% BSA in PBS. Click-iT reaction cocktail was prepared as in the protocol and added to the samples for 30 minutes at room temperature while rocking (protected from light). The reaction cocktail was removed and the samples were washed with 1mL of 3% BSA in PBS. The samples were then stained with DAPI according to the protocol and then stained with primary and secondary antibodies following the whole mount organoid immunofluorescence staining protocol as previously described. Samples were imaged using the Nikon confocal laser scanning microscopy.

## Data Analysis

### Statistical analysis

CD31 immunofluorescence staining was analyzed using Threshold on ImageJ (https://serc.carleton.edu/eet/measure_sat2/part_4.html). Images were converted into 8-bit and the Threshold dialog window was used to highlight CD31 positive pixels in the spheroids. Additionally, micrometers were selected under Analyze > Set Scale to measure the area in micrometers. The area of the spheroids was selected using the Freehand Selection tool in the ImageJ toolbar. By clicking Area and Limit to Threshold when setting measurements to analyze under the Analyze bar, we measured only the highlighted pixels within the spheroid area that was selected using the Freehand Selection tool. Positive percentage area was calculated using the positive threshold area divided by the area of the spheroid. Results were graphed using GraphPad and results were analyzed using one-way ANOVA.

## Acknowledgements

We thank all members of the Li laboratory for their insightful discussions and contributions to this work. We also acknowledge the Center for Biologic Imaging at the University of Pittsburgh for their support with spheroid imaging. We are grateful to Aidan Dorn for his assistance with imaging the spheroids. This research was supported by NIH grants R00HL133472 and DP2HL163745, the SVRF award from Additional Ventures, and the CMRF award from the University of Pittsburgh.

## Contributions

C.C., S.C., and G.L. designed and conducted the experiments. J.X. bred the mice. C.C. and G.L. drafted the manuscript, and all authors contributed to its editing.

## Supplemental Figure Legends

**Supplementary Figure 1:**
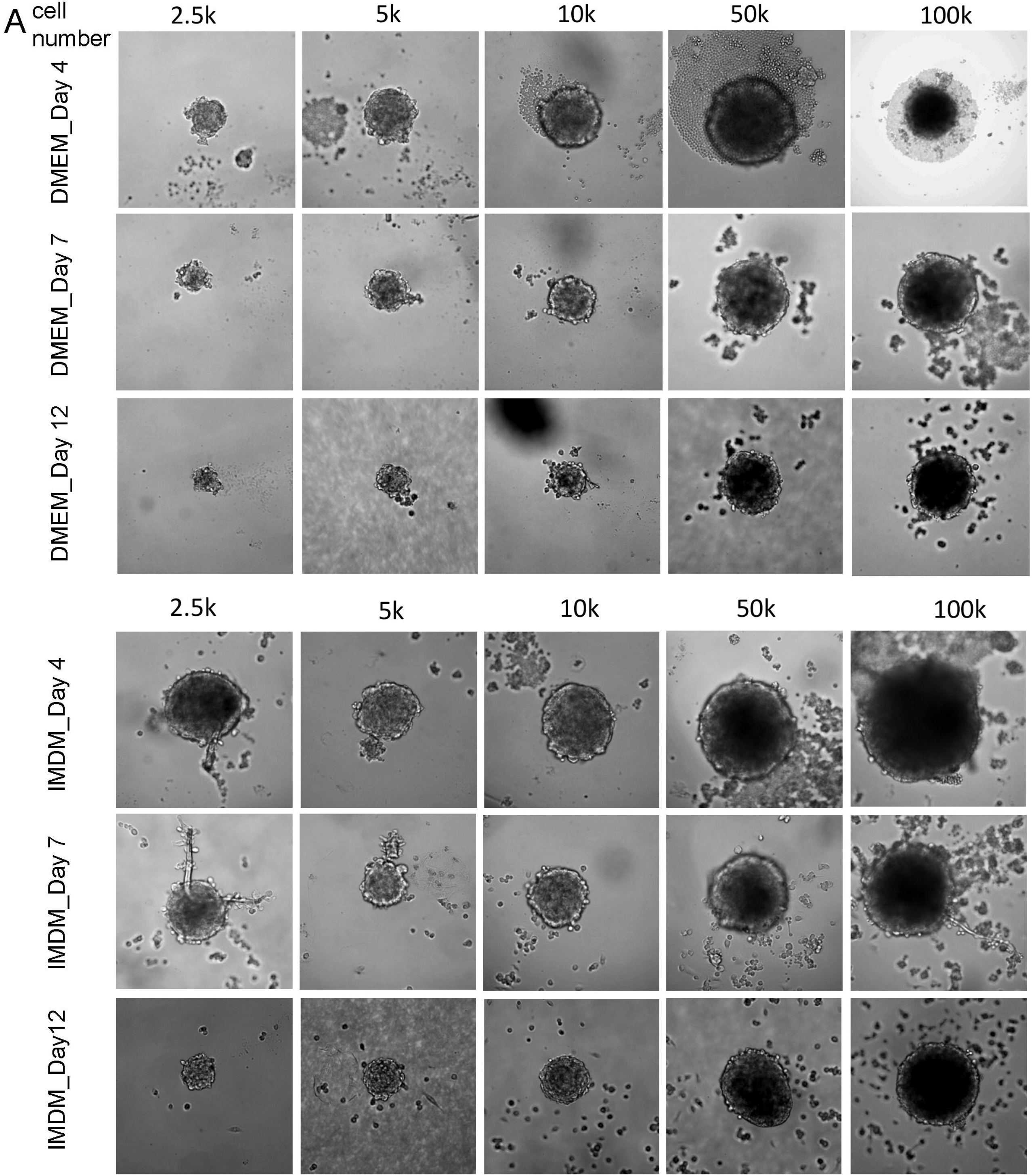
Brightfield images of ventricular spheroids with different cell densities cultured in DMEM and IMDM-based medium.

**Supplementary Figure 2:**
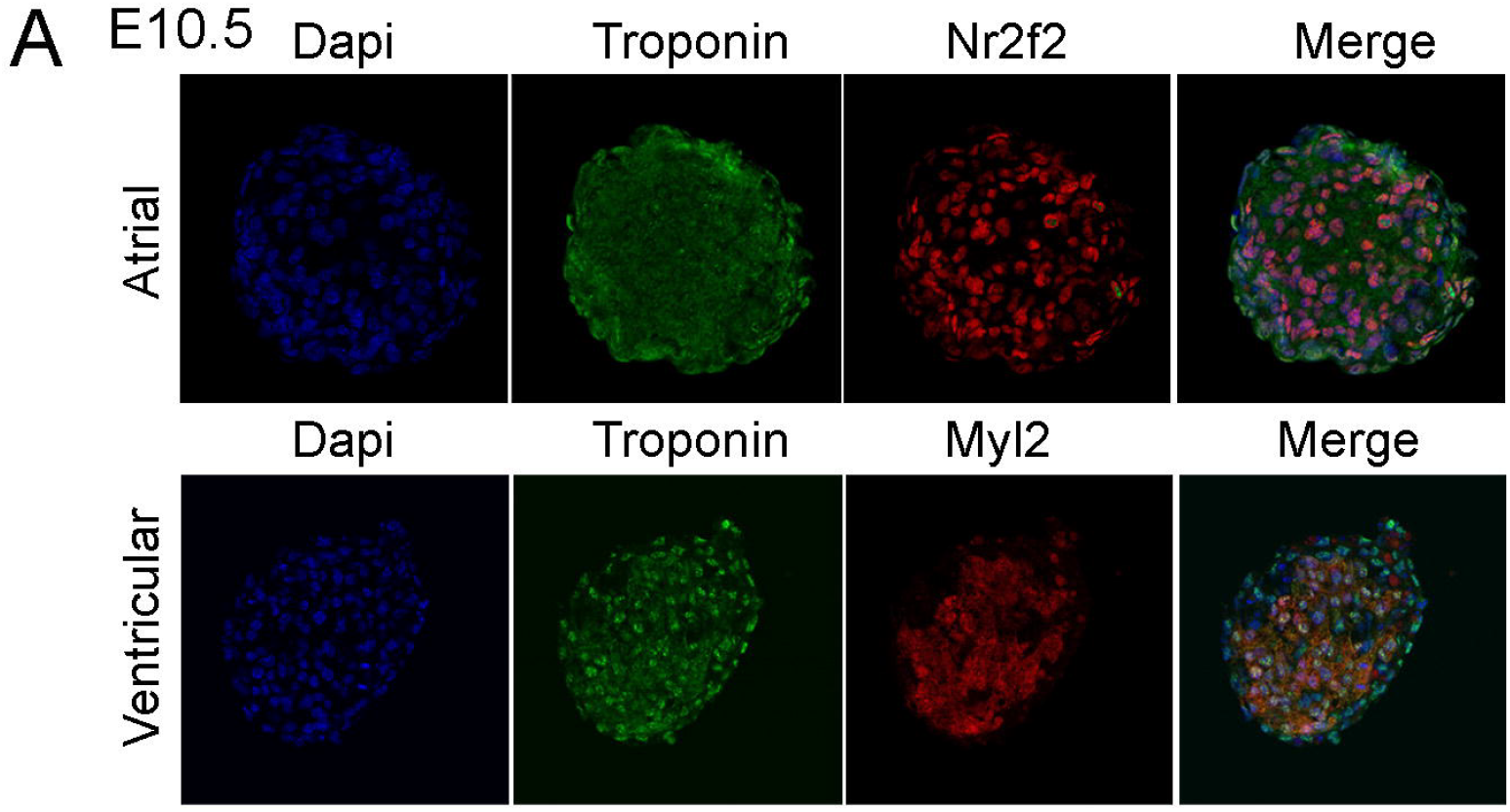
IF staining of atrial and ventricular CM lineage genes in atrial and ventricular spheroids generated with E10.5 heart cells.

**Supplementary Figure 3:**
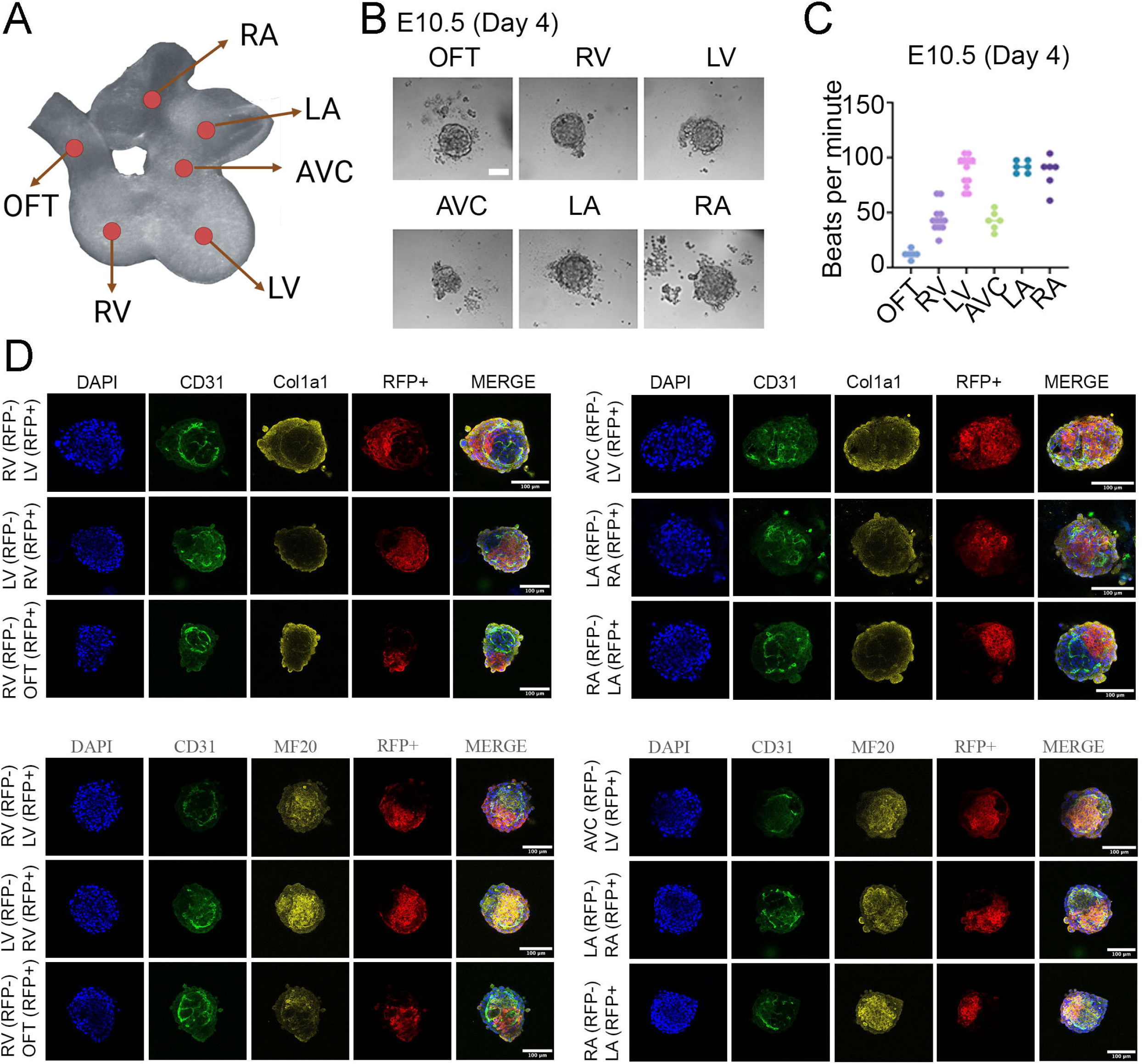
Generation of cardiac region-specific spheroids and fusions. (A) Schematic of the six heart regions in E10.5 mouse hearts: outflow tract (OFT), right ventricular (RV), left ventricular (LV), atrioventricular canal (AVC), left atrial (LA), and right atrial (RA). (B) Representative brightfield images of the six-heart regionderived spheroids. Scale bar =100 μm. (C) Beating rates of region-specific spheroids. (D) IF images of the fused spheroids from adjacent regions. scale bar=100 μm.

**Supplementary Figure 4:**
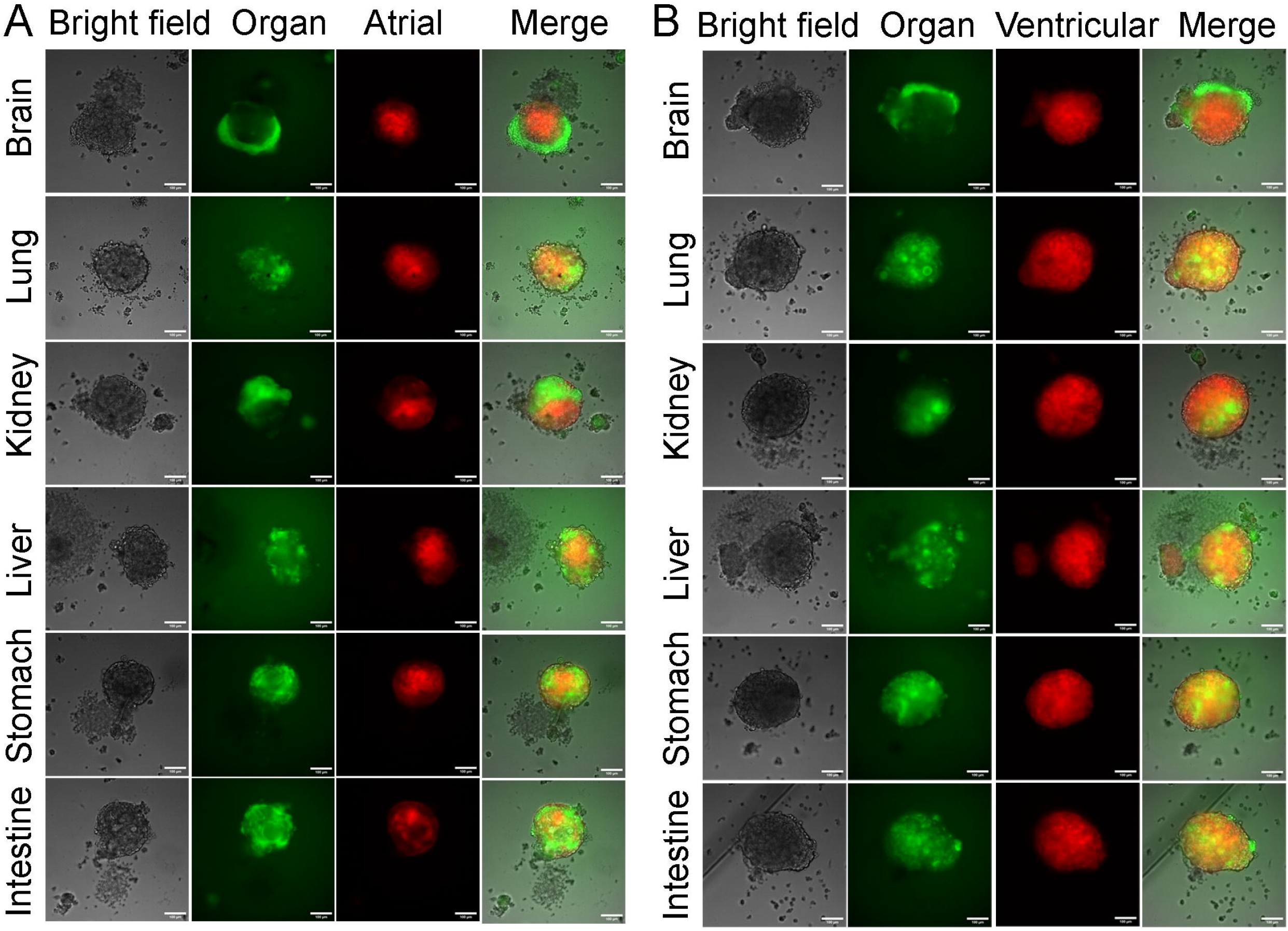
Spheroids generated by mixing atrial or ventricular cells (RFP+) with different organ cells (GFP+). Scale bar=100 μm.

